# *Wolbachia* effects on thermal preference of natural *Drosophila melanogaster* are influenced by host genetic background, *Wolbachia* type and bacterial titer

**DOI:** 10.1101/2023.07.31.551304

**Authors:** Anton Strunov, Charlotte Schoenherr, Martin Kapun

**Affiliations:** Center for Anatomy and Cell Biology, Medical University of Vienna, Austria; Central Research Laboratories, Natural History Museum of Vienna, Austria

**Keywords:** *Drosophila*, *Wolbachia*, titer, Tp, qPCR, environment

## Abstract

Temperature plays a fundamental role for the fitness of all organisms. In particular, it strongly affects metabolism and reproduction in ectotherms that have limited physiological capabilities to regulate their body temperature. Ectotherms thus have to maintain thermal homeostasis by behavioral adjustments. The influence of temperature variation on the physiology and behavior of ectotherms is well studied but we still know little about the influence of symbiotic interactions on thermal preference (*T*_p_) of the host. The *Wolbachia*-*Drosophila* host-symbiont system represents an ideal model for addressing these questions. A growing number of studies demonstrated that different *Wolbachia* types can influence *T*_p_ in different *Drosophila* species, but these results may be confounded by the use of long-term *Drosophila* lab-strains that may not be representative for natural fly populations. To account for this, we investigated the effect of *Wolbachia* on *T*_p_ in wild-type *D. melanogaster* flies recently collected from nature. Consistent with previous data, we found reduced *T*_p_ compared to an uninfected control in one of two fly strains infected with the wMelCS *Wolbachia* type. Additionally, we, for the first time, found that *Wolbachia* titer variation influences thermal preference of the host fly. These data indicate that the interaction of *Wolbachia* and *Drosophila* resulting in behavioral variation is complex and strongly influenced by the genetic background of host and symbiont. Our results further emphasize the necessity for more in-depth studies to better understand the evolutionary significance of *T*_p_ variation influenced by *Wolbachia* in natural *Drosophila* populations.

## Introduction

Temperature is a key environmental factor that has a profound impact on the physiology, reproductive success and fitness of all organisms on our planet (Angilletta *et al*., 2002; Johnston and Bennett, 2008; Hoffmann, 2010). In particular, ectotherms with restricted physiological capabilities to regulate their body temperature are challenged by fluctuating ambient temperatures (Huey and Berrigan, 2001; Angilletta *et al*., 2002; Hoffmann *et al*., 2003; Angilletta, 2004).

However, ectothermal organisms often account for environmental temperature variation by behavioral adjustments to avoid harmful or potentially lethal extreme conditions and to keep their body within an optimal physiological temperature range. Thermal preference (*T*_p_), is thus defined as the narrow thermal window that organisms choose when exposed to a broad range of different temperatures. Thus, this preferred temperature often corresponds to thermal conditions at which behavioral, physiological and other traits are optimized (Dillon *et al*., 2009). However, *T*_p_ is not absolute for a given species and can be highly variable with respect to the geographic origin of an organism. For example, both *Drosophila melanogaster* (Rajpurohit *et al*., 2017) and *D. subobscura* (Castañeda *et al*., 2013) from higher latitudes prefer lower mean temperatures compared to low-latitude flies which indicates that variation in *T*_p_ can have an adaptive value. Nevertheless, *T*_p_ is generally not genetically hard-wired but can be highly plastic, depending on life stage, physiology and other abiotic and biotic ecological factors (Dillon *et al*., 2009; Strunov, Schoenherr, *et al*., 2023).

Symbionts, ranging from mutualists to parasites, may further influence their hostś *T*_p_ which can lead to complex eco-evolutionary dynamics (de Roode and Lefèvre, 2012; Żbikowska and Cichy, 2015; Corbin *et al*., 2017). Ectothermal hosts, for example, can shift their *T*_p_ to higher temperatures (“behavioral fever”; Kluger, 1979; Thomas and Blanford, 2003) or lower temperatures (“behavioral anapyrexia”; Müller and Paul, 1993; Moore and Freehling, 2002; Fedorka *et al*., 2016) as a form of self-medication by moving to thermal environments outside the optimum of their parasitic symbiont.

Despite a growing body of literature reporting variation in host behavior influenced by symbionts, we still poorly understand functional links and the evolutionary impact of thermal preference dynamics in host-symbiont associations. The fruit fly *D. melanogaster* and the gram-negative bacteria *Wolbachia* represent a well-studied host-symbiont system that has proved eminently suitable to address these unresolved questions (Trinder *et al*., 2017). *D. melanogaster* is distributed on all continents except Antarctica (Keller, 2007; Haudry *et al*. 2020) and thus characterized by well-documented and widespread adaptations to temperate and tropical environments (Hoffmann *et al*., 2003). Moreover, it is one of the oldest and best-studied genetic model organisms (Schneider, 2000; Hales *et al*., 2015; Casillas and Barbadilla, 2017). The gram-negative bacterium *Wolbachia* is a facultative endosymbiont from the order rickettsiales, which is found in more than 40% of all terrestrial arthropod taxa (Zug and Hammerstein, 2012). *Wolbachia* is a well-studied, long-standing bacterial model because of its fundamental effects on host reproduction, physiology and immunity (Fry and Rand, 2002; Fry *et al*., 2004; Werren *et al*., 2008; Miller, 2013). It can act as a reproductive parasite that enhances its own transmission through manipulations of the host reproduction which may include male killing, feminization, parthenogenesis and cytoplasmic incompatibility (reviewed in Kaur *et al*., 2021). However, *Wolbachia* can also exhibit mutualistic effects by providing antiviral protection (Teixeira *et al*., 2008; Moreira *et al*., 2009; Osborne *et al*., 2009) and by increasing host fecundity (Miller *et al*., 2010; Strunov *et al*., 2022).

In addition, a handful of recent studies indicate that *Wolbachia* can manipulate thermal behaviour of *D. melanogaster*. Two studies reported that flies infected with *Wolbachia* tend to prefer cooler temperatures than uninfected controls (Arnold *et al*., 2019; Truitt *et al*., 2019). Particularly the *Wolbachia* type wMelCS, which is the second most common *Wolbachia* type in natural *D. melanogaster* (Riegler *et al*., 2005, 2012), was found to have a significantly stronger negative effect on *T*_p_ than the most common wMel type or uninfected controls. Truitt et al. (2019) speculated that a reduced *T*_p_ of flies infected with wMelCS might negatively influence the fitness of infected flies, e.g., due to slower development in colder environments compared to flies infected with wMel that prefer higher temperatures (Strunov *et al*., 2022). Such negative fitness effects could have contributed to the recent and rapid replacement of the ancestral wMelCS type by the nowadays dominant wMel type in worldwide *D. melanogaster* populations (Riegler *et al*., 2005; Strunov, Kirchner, *et al*., 2023).

However, two recent follow-up studies that further investigated the influence of *Wolbachia* on host *T*_p_ failed to identify similar shifts towards lower *T*_p_’s in flies infected with wMelCS (Hague *et al*., 2020; Strunov, Schoenherr, *et al*., 2023). Such inconsistencies indicate that the influence of *Wolbachia* on the host *T*_p_ may be subtle and strongly influenced by environmental factors, the host physiology and the host genetic background (Strunov, Schoenherr, *et al*., 2023). Notably, all of the four aforementioned studies were based on experiments using long-standing *Drosophila* lab-strains. Maintenance of flies in the lab with an excess of food, constant inbreeding, and relaxation from typical environmental constraints, potentially introduce artefacts in behavioral and physiological traits (Markow, 2015). It thus remains unknown to which extent mutation accumulation and lab-adaptation may confound the interpretation of these data and may preclude drawing conclusions about the evolutionary relevance of these findings in natural populations.

Moreover, the mechanisms that may contribute to behavioral differences among hosts carrying wMel or wMelCS remain unknown. Chrostek et al. (2013) first reported that *Wolbachia* titer levels in wMelCS-infected flies were significantly higher than in wMel-infected hosts. Elevated titers of wMelCS may lead to increased physiological stress to the host and could thus modulate behavioral responses that shift the *T*_p_ towards colder temperatures to alleviate fitness costs from high wMelCS titers by reducing the bacterial proliferation rates in colder temperatures. Inconsistent results in the four aforementioned studies could thus potentially stem from titer variation due to differences in host strains and in the experimental setups (Strunov, Schoenherr, *et al*., 2023).

To address these unresolved questions, we focused on wild-type fly strains infected with different *Wolbachia* types that were only recently collected from natural populations at the endpoints of a latitudinal gradient in Europe. We investigated the effects of *Wolbachia* infections on host *T*_p_ by comparing thermal preference of naturally infected flies to subsets of the same fly strains that we cured from *Wolbachia* infections by antibiotics treatment. Furthermore, we, for the first time, tested for a direct effect of *Wolbachia* titer on thermal preference using quantitative PCR (qPCR). The central goal of our study was thus to better understand the influence of *Wolbachia* on thermal behavior in natural *Drosophila* populations and to elucidate the mechanisms that may trigger behavioral responses in the *Drosophila*-*Wolbachia* symbiotic system.

## Materials & Methods

### Drosophila melanogaster isofemale strains and their infection status

For our analyses, we focused on eight isofemale *Drosophila melanogaster* strains collected in October 2018 from wild populations in Recarei/Portugal and Akka/Finland (**Table 1**). We first assessed their infection status using *Wolbachia*-specific *wsp* primers (Zhou *et al*., 1998). For six strains, which were positively tested for *Wolbachia* infections, we further inferred the *Wolbachia* type based on PCR with primers for the VNTR-141 locus, which is diagnostic for the wMel and wMelCS *Wolbachia* types (Riegler *et al*., 2012). To obtain uninfected controls for each of the eight strains, we treated a subset of each strain with antibiotics (AB; 0.1% Rifampicin) for three generations to eliminate bacterial infections, and subsequently restored the microbial gut flora (GFR) by placing the AB-treated flies in food vials with freshly deposited fly faeces from untreated males of the same strain for two generations. Fly stocks were maintained in incubators at 24°C and 60% humidity with a 12h:12h light/dark cycle prior to experiments.

**Table 1.** Description of wild-type and lab strains of *D. melanogaster* used in the present study.

| Sample ID | Date of sampling | Sampling location | Time of Tp experiment | *Wolbachia* status |
| --- | --- | --- | --- | --- |
| Uninf_Po_1<br>Uninf_AB_Po_1 | 17.10.19 | Recarei, Portugal | Sep. 2020 | w-AB treatment |
| Uninf_Po_2<br>Uninf_AB_Po_2 | 18.10.19 | Recarei, Portugal | Mar. 2021 | w-AB treatment |
| wMel_Po_1+<br>wMel_Po_1- | 10.01.19 | Recarei, Portugal | Sep. 2020 | wMel<br>AB treatment |
| wMel_Po_2+<br>wMel_Po_2- | 10.06.19 | Recarei, Portugal | Mar. 2021 | wMel<br>AB treatment |
| wMel_Fi_1+<br>wMel_Fi_1- | 10.07.19 | Akka, Finland | Mar. 2021 | wMel<br>AB treatment |
| wMel_Fi_2+<br>wMel_Fi_2- | 10.09.19 | Akka, Finland | Mar. 2021 | wMel<br>AB treatment |
| wMelCS_Po_1+<br>wMelCS_Po_1- | 10.10.19 | Recarei, Portugal | Sep. 2020 | wMelCS<br>AB treatment |
| wMelCS_Po_2+<br>wMelCS_Po_2- | 10.03.19 | Recarei, Portugal | Mar. 2021 | wMelCS<br>AB treatment |

### Temperature preference measurement

For *T*_p_ measurements in *Drosophila* adults, we used a thermal gradient apparatus with two narrow arenas (Strunov, Schoenherr, *et al*., 2023). In short, the device consisted of an elongated aluminium plate with a plexiglass cover that contained a central hole to apply test subjects. Thin plexiglass dividers formed two narrow arenas with 6 mm diameter between the two plates. The length of the arena was restricted to 30 cm by applying foam stoppers at the ends of each lane to prevent flies from escaping and to allow a constant airflow. Moisturised Whatman paper was used as a bottom layer to cover the aluminium plate in the arenas and to prevent the test subjects from desiccation. After each experiment, we exchanged the Whatman paper and rinsed all parts of the apparatus with soap and hot water to remove any potential olfactory contaminants, which might interfere with thermal preference behaviour of flies.

To establish a constant temperature gradient, we attached two peltier elements, that were placed below the aluminium plate at the endpoints of the prospective temperature gradient, to power supplies set to 2 V and 3 V for cooling and heating, respectively. Temperature was constantly monitored at both ends and in the middle of the arena with digital temperature sensors (Analog devices, USA), starting 20 min prior to the onset of each experiment. The data from the thermal sensors were collected with a custom python script on a Raspberry pi 3B+ computer (Raspberry Pi Foundation, UK) in 10 sec intervals and saved as text files (see https://github.com/capoony/DrosophilaThermalPreference).

All experiments were carried out on two thermal gradient devices simultaneously in a dark isolated room equipped with air-conditioning to keep the ambient temperature stable at 21-23°C, which was constantly monitored with a data logger. The position of the flies within the devices were captured by an infrared camera attached atop the two thermal gradient devices. Images were taken every 30 seconds and stored on the Raspberry pi computer for each run (Strunov, Schoenherr, *et al*., 2023). The coordinates of each individual fly and of the thermal sensors were manually assessed in ImageJ (Schneider et al., 2012) based on infrared images captured 20 minutes after the onset of each experiment. Following the approach described in Strunov, Schoenherr et al. (2023), we used a custom python script to estimate *T*_p_ for every fly using information of its position relative to the linear temperature gradient between the temperature measurement points.

Prior to the experiments, we selected cohorts of 20 females and 20 males, which were one week old (± two days). We transferred the flies to fresh glass vials with medium by gently applying CO_2_ and then allowed the flies to recover for two days. Each experimental run included untreated and AB-treated flies of a given strain placed in neighbouring arenas to keep uncontrolled environmental influences consistent when testing for the effects of *Wolbachia* on *T*_p_ in flies of the same genetic background. Prior to each experiment, we randomly assigned experimental cohorts to the two devices that operated simultaneously. Flies were then gently collected from their vial with a handmade fly aspirator and transferred to the thermal gradient devices through holes in the middle of the plexiglass covers. Then, the holes were tightly sealed with cotton, the light was turned off and measurements were taken during a 20-30 minutes period. After the experiment was finished, flies were anaesthetized with CO_2_ within the apparatus, collected into 1.5ml tubes and frozen at -20°C for further analyses (see below).

Since time of the day can have a strong influence on *T*_p_ in *Drosophila* (Chen *et al*., 2022), we performed a maximum of four experiments per day to avoid potential bias from behavioral changes due to circadian rhythms. The time of the runs during the day was usually as follows: first two runs were carried out from 10 am to 12 am, last two runs from 1 pm to 3.30 pm. To quantify *T*_p_ of individual flies in the arena, we used the images of fly locations taken 20 min after the onset of the experiment similar to Truitt et al. (2019) and Strunov, Schoenherr et al. (2023).

### qPCR-based analysis of relative Wolbachia titers

We used quantitative PCR (qPCR) following the approach in Hague et al. (2020) to assess relative *Wolbachia* titers from whole-body DNA extracted with the Gentra Puregen DNA extraction kit (Qiagen). We compared cycle threshold (*Ct*) values of the single-copy *RPL32* gene of *Drosophila melanogaster* (primer forward - 5’-CCGCTTCAAGGGACAGTATC-3’, primer reverse - 5’-CAATCTCCTTGCGCTTCTTG-3’) and a single copy ankyrin repeat gene *WD0550* of *Wolbachia* (primer forward - 5’-CAGGAGTTGCTGTGGGTATATTAGC-3’, primer reverse - 5’-TGCAGGTAATGCAGTAGCGTAAA-3’). PCR reactions were carried out on a CFX384 Real-Time PCR detection system (Bio-Rad) in three-fold technical replication for each sample. All samples were randomly distributed on 384-well plates to exclude possible plate-specific batch effects. Before setting up the main experiments, we set up dilution series to produce efficiency curves for each primer pair to confirm that the primer efficiency was close to 100%. Each of the PCRs was performed in a 10 µl reaction volume consisting of 5 μl Luna Mix (NEB), 0.5 μl of each primer (3.6 mM), and 4 μl of DNA (diluted to 0.5 ng/μl). The following thermal cycling protocol was used for amplification: initial 5 min at 50°C, denaturation for 5 min at 95°C, followed by 40 cycles of 30 s at 95°C, 45 sec at 60°C, and 5 s at 65°C. Cycle threshold (*Ct*) values were obtained using Bio-Rad CFX Manager software with default threshold settings ignoring *Ct* values > 30. To preclude inconsistent results, we discarded and repeated PCRs for samples whose standard deviations between technical replicates exceeded 0.5.

To calculate relative *Wolbachia* titers, we followed the approach in Hague et al. (2020) using the Pfaffl method (Pfaffl, 2001) by standardising bacterial titers based on *Ct* values of the host *RPL32* gene as shown in formula (1), where Ct is the average *Ct* value across three technical replicates for the *Wolbachia* and the host gene, respectively.

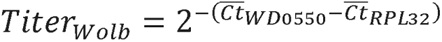

### Statistical analyses

We tested for the effects of *Wolbachia* infection type, AB-treatment, geographic origin and other environmental factors on *T*_p_ of the *D. melanogaster* host using linear mixed models (LMMs) in *R* (R Core Team, 2019) based on the “lmer” function in the lme4 package (Bates *et al*., 2015). thermal preference of every individual was considered the dependent variable, and *infection type*, *AB-treatment* and *geographic origin* as the fixed factors. Moreover, we always included the factor *replicate experiment ID* nested within *sample ID* as a random factor in each of our models. We tested for significance of the fixed factors and all possible interactions in the *T*_p_ experiments using the *anova* function in *R* and calculated a Type-III analysis of deviance comparing nested models. Whenever a factor with more than two factor levels had a significant effect, we performed post-hoc tests using Tukey’s HSD method as implemented in the *emmeans* package (Lenth *et al*., 2019). Additionally, we tested for associations between relative *Wolbachia* titers and *T*_p_ in each investigated strain separately using linear models and tested for significance using the summary function in *R*. All *R* code can be found at https://github.com/capoony/DrosoNatThermPref.

## Results and Discussion

In this study, we investigated the influence of *Wolbachia* infections on thermal behavior in naturally infected wild-type *Drosophila melanogaster* from two populations at the endpoints of a latitudinal temperature gradient in Europe. We used a custom-built thermal gradient device (Strunov, Schoenherr, *et al*., 2023) to assess individual thermal preference (*T*_p_) in eight wild-caught isofemale strains of *D. melanogaster* collected in Recarei, Portugal and Akka, Finland in 2019 (**Table 1**). A subset of each line was cured from *Wolbachia* infections by antibiotic treatment followed by three generations of gut flora restoration to generate uninfected controls (see Materials and Methods). In total, we analyzed four strains infected with the *w*Mel *Wolbachia* type, two strains infected with *w*MelCS, and two naturally uninfected strains and compared these to their antibiotics-treated counterparts of each strain. Our experiments were carried out in 10-13 biological replicates per isofemale strain and treatment with a total of 6,182 flies investigated.

### T_p_ is influenced by Wolbachia infections and humidity but not geographic origin

Irrespective of AB-treatment we observed highly significant differences in thermal preference among all isofemale lines (**Figure 1**; LMM; Χ^2^=46.6, df=7, *p* < 0.001; **Table 2**). Mean *T*_p_ ranged from 20°C in a naturally uninfected strain from Portugal (Uninf_Po_2; **Table S1**) to 22.9°C in a Finish strain infected with *w*Mel (wMel_Fi_2; **Table S1**).

**Figure 1.**
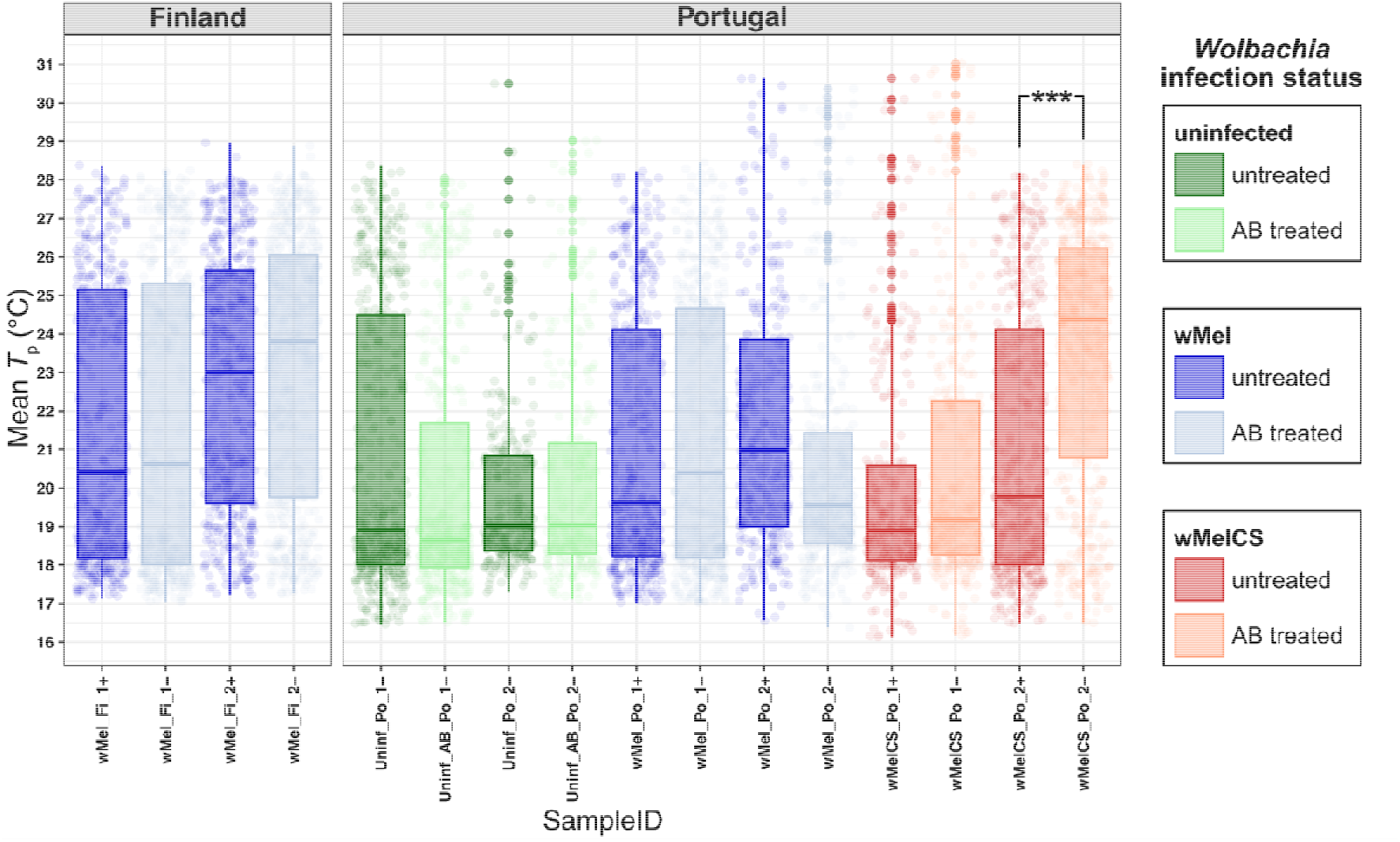
Thermal preference (*T*_p_) of wild-type *D. melanogaster* flies from Portugal and Finland in the presence and absence of *Wolbachia* infection. Boxplots are showing *T*_p_ in eight natural *Drosophila* strains from Finland and Portugal that were either naturally uninfected (green), or either naturally infected with wMel (blue) or wMelCS (red). The “-” symbol in the sample names and light colors of the boxplots depict strains that underwent antibiotic treatment and subsequent restoration of the gut flora; The “***” symbol indicates a *p*-value < 0.0001.

**Table 2.** Linear mixed model testing for the effects of sample ID, AB treatment, ambient temperature, humidity and the interaction between sample ID and AB treatment on *T*_p_. The columns show the factor names, the Χ values, the degrees of freedom and the corresponding *p*-values inferred by analyses of deviance, respectively. Significant results are highlighted in bold.

| *Factor* | $\chi^2$ | *df* | $Pr(>\chi^2)$ |
| --- | --- | --- | --- |
| **ID** | **46.6** | **7** | **1.00E-07** |
| AB treatment | 2.7 | 1 | 0.098 |
| ambient temperature | 2.3 | 1 | 0.128 |
| **humidity** | **27.2** | **1** | **2.00E-07** |
| ID x AB treatment | 11.7 | 7 | 0.112 |

Furthermore, we found no difference in *T*_p_ with respect to antibiotics treatment in either of the two naturally uninfected strains (Tukey-HSD; *z*-ratios = 0.49 and - 0.27; *p* > 0.05 for Uninf_Po_1 and Uninf_Po_2, respectively; **Table 3**), which served as controls. We thus conclude that potential perturbations of the gut microbiome from the AB-treatment itself did not influence average thermal preference. In only one of the isofemale strains (wMelCS_Po_2) we identified significant differences in thermal preference between untreated and AB-treated flies (mean Δ*T*_p_=2.4°C; Tukey-HSD; z-ratio = -3.34; *p* < 0.001; **Table 3**), which indicates that the infection with *w*MelCS reduced thermal preference in this isofemale line similar to previous data (Arnold *et al*., 2019; Truitt *et al*., 2019).

**Table 3.** Tukey HSD posthoc tests, testing for the effect of antibiotics treatment on *T*_p_ in different host.

| *contrast (AB-treatment)* | *strain* | *z-ratio* | *p-value* |
| --- | --- | --- | --- |
| no - yes | wMel_Fi_1 | -0.484 | 0.629 |
| no - yes | wMel_Fi_2 | -0.560 | 0.575 |
| no - yes | wMel_Po_1 | -0.642 | 0.521 |
| **no - yes** | **wMelCS_Po_2** | **-3.340** | **0.001** |
| no - yes | Uninf_Po_1 | 0.493 | 0.622 |
| no - yes | Uninf_Po_2 | -0.275 | 0.783 |
| no - yes | wMelCS_Po_1 | -0.567 | 0.571 |
| no - yes | wMel_Po_2 | 0.873 | 0.383 |

Next, we specifically tested for significant *T*_p_ differences between treated and untreated strains with respect to *Wolbachia* infection type and for differences among strains according to geographic origin. While we neither observed differences in *T*_p_ according to geographic origin alone (LMM; Χ^2^=0.3, df=1, *p*=0.568; **Table S2**) nor interactions among geographic origin and AB treatment (LMM; Χ^2^=0.4, df=1, *p*= 0.518; **Table S2**), we identified a significant interaction between *Wolbachia* infection type and AB treatment (LMM; Χ^2^=7.0, df=2, *p*=0.031; **Table S2**). Similar to the analysis above, a follow-up post-hoc test revealed that only the difference in *T*_p_ between AB-treated and untreated flies of the isofemale line wMelCS_Po_2 was significant (mean Δ*T*_p_=2.4°C; Tukey-HSD; z-ratio = -2.6; p-value < 0.01; **Table S3**).

In summary, our results indicate that the genetic background of an isofemale strain has a strong influence on thermal preference, which is in line with previous studies (Hague *et al*., 2020; Serga *et al*., 2021; Strunov, Schoenherr, *et al*., 2023). We nevertheless found that *w*MelCS infections can have a negative influence on thermal preference, which was shown previously (Arnold *et al*., 2019; Truitt *et al*., 2019; but see Hague *et al*., 2020; Strunov, Schoenherr, *et al*., 2023). Since we observe that only one line naturally infected with *w*MelCS exhibits an increased thermal preference when treated with antibiotics, we speculate that the influence of *Wolbachia* infections on thermal preference may often be subtle and strongly influenced by the host genetic background and potentially also by interactions with other environmental factors. Consistent with previous data (Strunov, Schoenherr, *et al*., 2023), we observed that the amount of humidity (LMM; Χ^2^=27.2, df=1, *p* < 0.001; **Table 2**), but not ambient temperature (LMM; Χ^2^=2.3, df=1, *p*=0.128; **Table 2**) had a highly significant influence on thermal behaviour. Subtle behavioural effects of bacterial infection masked by a strong influence of host genetic background and environmental variation may thus explain why the effects of *Wolbachia* on thermal preference were not consistently identified in other studies (see, for example, Hague *et al*., 2020; Strunov, Schoenherr, *et al*., 2023).

In contrast to Rajpurohit et al. (2017) who reported a positive correlation between *T*_p_ and latitude along the North American east coast, we failed to identify thermal differences between strains from the endpoints of a latitudinal temperature gradient in Europe. However, we caution that the low number of isofemale strains from each of the two populations used in this study may confound a robust interpretation of our results in the context of spatial varying selection on thermal behaviour. Future studies with a stronger focus on population-wide effects are therefore needed to quantitatively address this specific question.

### T_p_ is influenced by Wolbachia titer variation and environmental cues

When investigating the consistency of thermal preference within strains and treatment, we observed substantial *T*_p_ variation among replicate experiments of the thermal preference assays (standard deviation from 2.4°C to 3.7°C depending on the line studied, **Table S1**). This further indicates that differences among the test subjects or differences in environmental conditions may contribute to variation in behavioural responses among the replicated experiments.

To investigate possible sources of variation among replicates, we tested if differences in *Wolbachia* titer contribute to thermal behaviour. Consistent with the hypothesis that higher *Wolbachia* titer is costly to the host and may thus trigger a shift towards lower *T*_p_ to reduce proliferation rates of the symbionts (Truitt et al., 2019), we found a highly significant negative correlation between mean *T*_p_ and mean bacterial titer in wMelCS_Po_2 (LM, *R*^2^=0.62, *t*=-5.675, *p*<0.001; **Figure 2**), which is naturally infected with *w*MelCS and exhibited lower *T*_p_ in infected flies compared to their AB-treated counterparts (**Figure 1**, **Table 3**).

**Figure 2.**
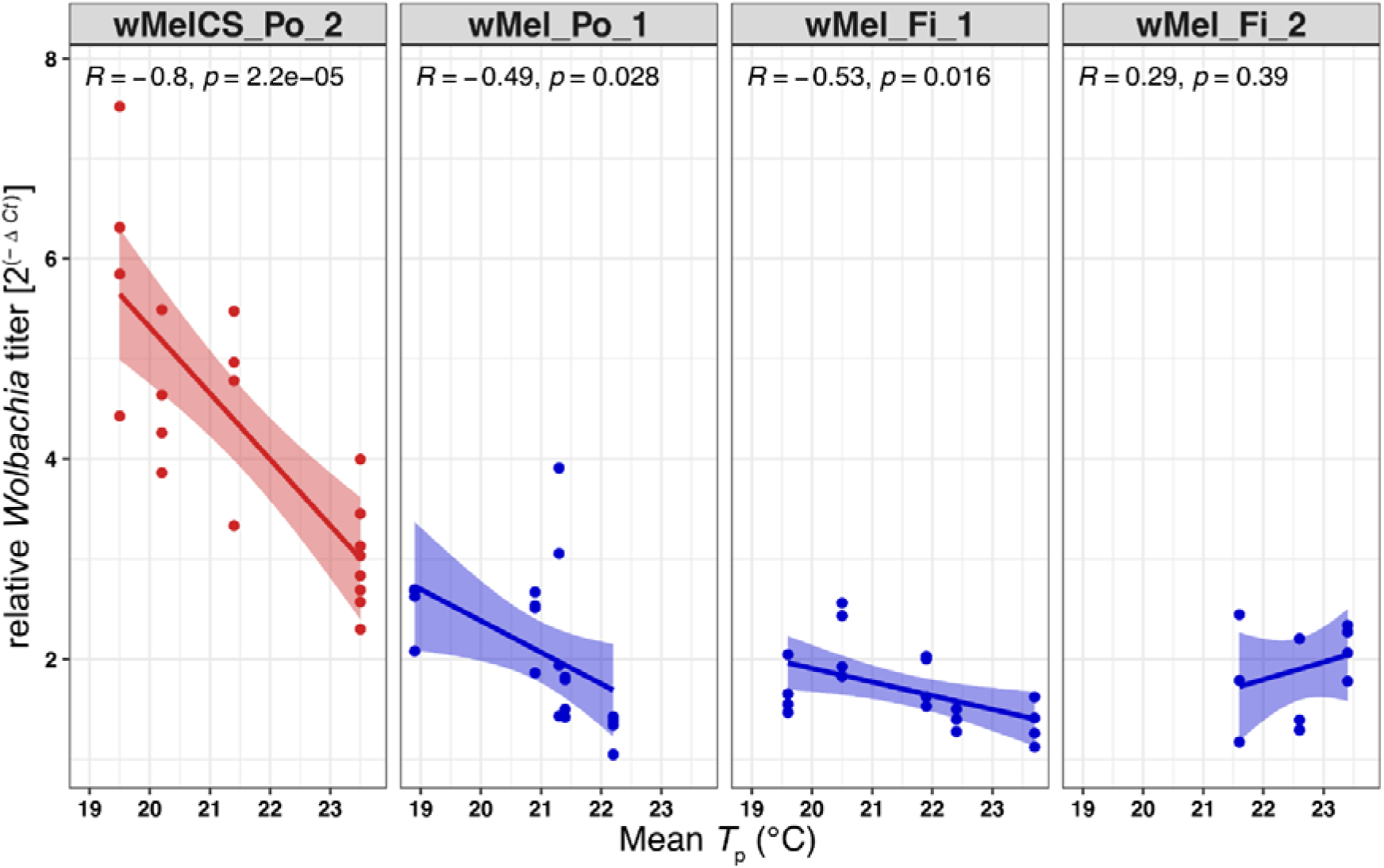
*Wolbachia* titer negatively influences host *T*_p_. Linear regression plot show significant negative correlations between *Wolbachia* titer (2^(-Δ*C*t)^) and mean preferred temperature (°C) for three *Drosophila* strains infected with wMelCS (red) or wMel (blue) but not for sample wMel_Fi_2 infected with wMel.

Surprisingly, two out of three wMel-infected strains from Portugal and Finland, which did not exhibit *T*_p_ difference with respect to their AB-treated controls (**Table S1**), also showed significant negative correlations between relative bacterial titer and thermal preference (wMel_Po_1: *R*^2^=0.24, *t*=-2.661, *p*=0.028; wMel_Fi_1: *R*^2^=0.28, *t*=-2.396, *p*=0.016; but see wMel_Fi_2: *R*^2^=0.02, *t*=-2.396, *p*=0.39; **Figure 2, Table S4**).

Unfortunately, we were not able to obtain data on bacterial titers of two additional strains with *w*MelCS and *w*Mel infections (wMelCS_Po_1 and wMel_Po_2, respectively), which were part of the aforementioned thermal preference assays. Given that flies of the second *w*MelCS-infected strain (wMelCS_Po_1) did not exhibit *T*_p_ differences compared to their AB-treated counterparts, we wanted to further test if bacterial titers in wMelCS_Po_1 were overall lower and more similar to *w*Mel-infected strains than wMelCS_Po_2. Such a pattern may explain why wMelCS_Po_1 did not show lower *T*_p_ when infected with wMelCS. We therefore assessed relative bacterial titers in laboratory stocks of the six naturally infected strains. Contrary to our hypothesis but consistent with literature (Chrostek *et al*., 2013), we found that both *w*MelCS-infected strains had significantly higher titer levels compared to all four *w*Mel-infected strains (Tukey-HSD; *p*-value < 0.0001; **Figure 3**; **Table S5**).

**Figure 3.**
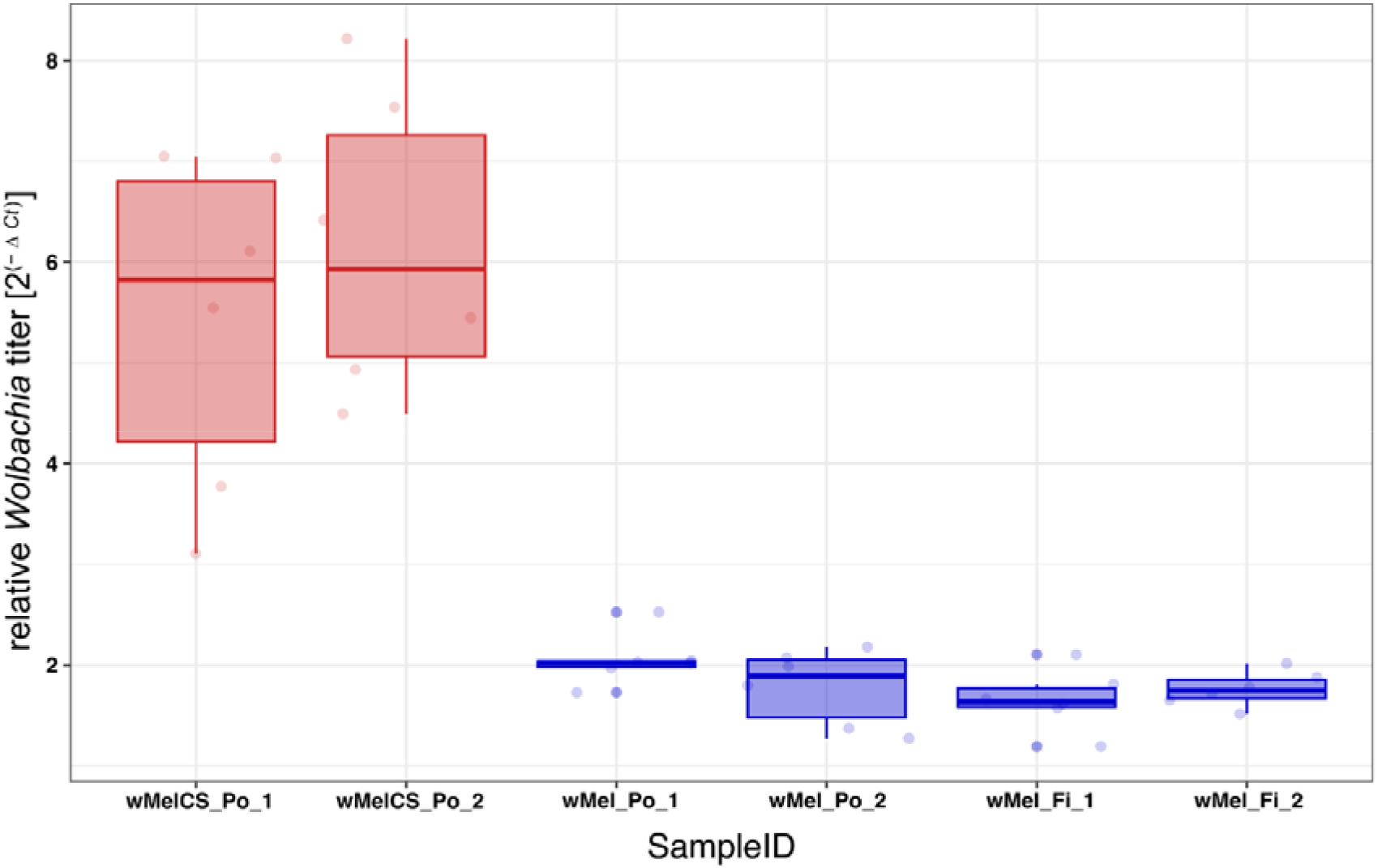
Relative *Wolbachia* titer in lab stocks of the six *D. melanogaster* strains used in this study. Note the significantly higher infection of wMelCS variant in comparison to wMel.

In summary, our results provide the first direct evidence that *Wolbachia* titers negatively influence thermal preference of its host. However, bacterial titer does not seem to be the only determinant of thermal preference, given that we do not observe consistent *T*_p_ differences among both wMelCS-infected strains and their AB-treated controls despite similarly high infection levels.

Other factors that may strongly influence the outcome of behavioral assays are social cues that can, for example, result in social crowding (Sayeed and Benzer, 1996; Dillon *et al*., 2009). In this study, we observed several replicated experiments where >80% of all tested flies were aggregating at the lower end of the thermal gradient (within 2°C from the minimum temperature; **Figure S1**). Since minimum temperatures along the temperature gradient never went below 16°C, a temperature where *D. melanogaster* can still successfully reproduce (Schnebel and Grossfield, 1986) and move (Giraldo *et al*., 2019; Davis *et al*., 2021), it is unlikely that clustering was the result of reduced mobility of flies from cold temperature exposure (but see Hague *et al*., 2020). We speculate that this phenomenon resulted from some form of edge attraction (Dillon *et al*., 2009; Soibam *et al*., 2012) and social clustering (Rooke *et al*., 2020). Curiously, we did not observe similar clusters at the hot end of the thermal gradient, which indicates that the higher temperature induces stronger aversion behaviour than cold temperature (Simões *et al*., 2021). To rule out that such clustering caused a bias in our results, we removed the corresponding replicate experiments from the dataset, which reduced the number of investigated flies by 15.2% (5,241 flies). We obtained qualitatively similar results when repeating the above-mentioned statistical analyses on the reduced dataset (**Tables S6, Figure S2**), which further indicates that the *w*MelCS-specific effect on *T*_p_ as observed in strain wMelCS_Po_2 is not an experimental artefact.

## Conclusion

In this study, we for the first time investigated the influence of *Wolbachia* on thermal preference in wild-type *Drosophila* hosts that were recently collected from two populations at the endpoints of a latitudinal temperature gradient in Europe. Consistent with previous findings in long-standing laboratory fly strains, we found that *Wolbachia* infections can lower *T*_p_ in flies naturally infected with the wMelCS *Wolbachia* type (Arnold *et al*., 2019; Truitt *et al*., 2019). Given that these patterns were only observed in a single fly strain, we conclude that the host genetic background has a strong influence on the strength and patterns of behavioural responses in the host flies (Hague *et al*., 2020; Serga *et al*., 2021; Strunov, Schoenherr, *et al*., 2023). However, when specifically testing for the effects of titer levels on thermal preference, we for the first time show that even subtle variations in bacteria titer can substantially influence the thermal response of the host. Remarkably, this effect was even found in wMel-infected flies, which are characterized by substantially lower titers than flies infected with wMelCS (Chrostek *et al*., 2013). We thus conclude that titer variation, rather than *Wolbachia* type, has a strong influence on the behavioral response investigated here. However, given that we did not observe consistent behavioral patterns in the natural *Drosophila* strains investigated in this study, the evolutionary significance of behavioral responses to *Wolbachia* infections by the host under natural conditions thus remains elusive to date. Further studies are needed to investigate in more detail how *Wolbachia* titer levels influence the physiology and resilience of their host in the context of environmental variation (Strunov *et al*., 2022) and, most importantly, how symbiont proliferation rates themselves are influenced by the ambient temperature (e.g., Elliot *et al*., 2002; Chrostek *et al*., 2021). This is particularly crucial in the light of ongoing applied research that focuses on using *Wolbachia* as a pest control agent (Bourtzis, 2008; Moreira *et al*., 2009; Iturbe-Ormaetxe *et al*., 2011; Pagendam *et al*., 2020; Sullivan, 2020). The influence of temperature on *Wolbachia* titer levels and potential subsequent ecological feedback loops from behavioral adjustments of the host flies may affect the predictability of *Wolbachia* infections on viral resistance and cytoplasmic incompatibility in disease vectors.

## Supporting information

Supportinf Information - Figures

## Acknowledgements

We are especially grateful to Wolfgang Miller who helped with the design of the experiment, who provided lab space and contributed to critical discussions of this project. We are also indebted to Marcel Freund at the University of Zürich who helped with the design and crafted the thermal gradient devices used in this study. We are very grateful to Fabian Gstöttenmayr, Elina Koivisto, Jasmin Jester and Marlene Muelbauer who helped with fly maintenance and fly food cooking and to Elisabeth Haring for providing critical feedback to the manuscript. This project was funded by a standalone grant from the Austrian Science Fund (FWF P32275) to Martin Kapun.

## Authors contribution

Anton Strunov involved in conceptualization, investigation, data curation, visualization, formal analysis, validation, writing—original draft; Charlotte Schoenherr involved in investigation, data curation and writing—review & editing; Martin Kapun involved in conceptualization, formal analysis, visualization, supervision, funding acquisition, writing— review & editing and project administration.

## Supplementary Figures

**Figure S1.**
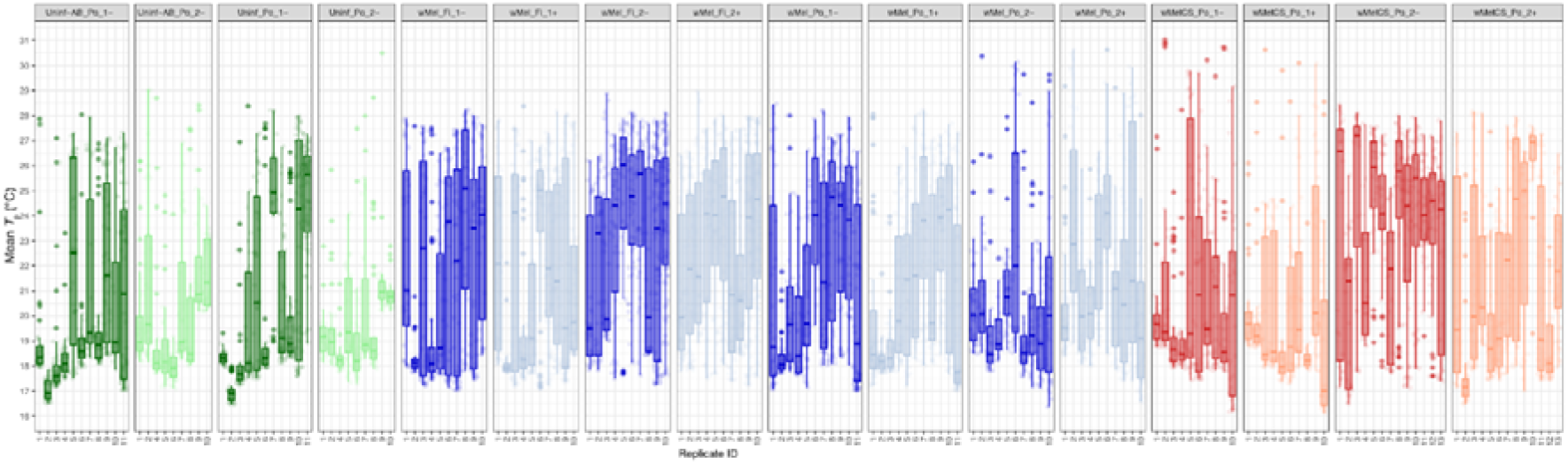
Boxplots showing thermal preference (*T*_p_) in replicated experiments using eight wild caught *Drosophila* strains that were either uninfected (green), infected with wMel (blue) or wMelCS (red). Light colors and the “-” symbol in the sample names indicate results for subsets of the strains treated with antibiotics. See Materials and Methods for more details.

**Figure S2.**
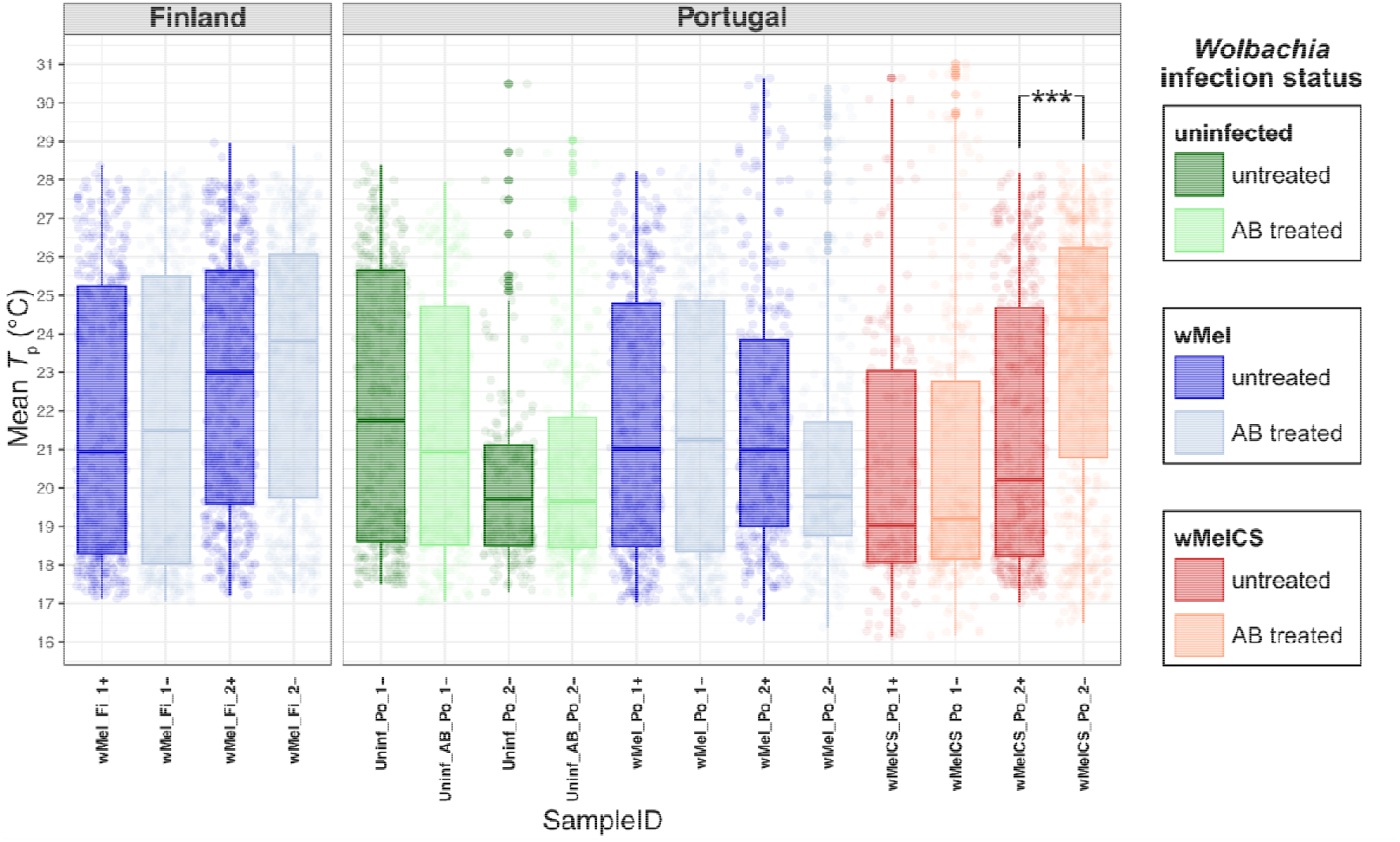
Similar to Figure 1, these boxplots are showing *T*_p_ in eight natural *Drosophila* strains from Finland and Portugal that were either naturally uninfected (green), or either naturally infected with wMel (blue) or wMelCS (red). The “-” symbol in the sample names and the light colors of the boxplots depict strains that underwent antibiotic treatment and subsequent restoration of the gut flora; In contrast to Figure 1, we here removed replicate experiments, where >80% of all flies cluster at the cold edge of the thermal gradient. The “***” symbol indicates a *p*-value < 0.0001.

