## Supplementary material for "*Wolbachia* effects on thermal preference of natural *Drosophila melanogaster* are influenced by host genetic background, *Wolbachia* type and bacterial titer": Supportinf Information - Figures

### Supplementary Figures:

##### **Figure S1.** Boxplots showing thermal preference (*T*_p_) in replicated experiments using eight wild caught *Drosophila* strains that were either uninfected (green), infected with wMel (blue) or wMelCS (red). Light colors and the “-” symbol in the sample names indicate results for subsets of the strains treated with antibiotics. See Materials and Methods for more details.

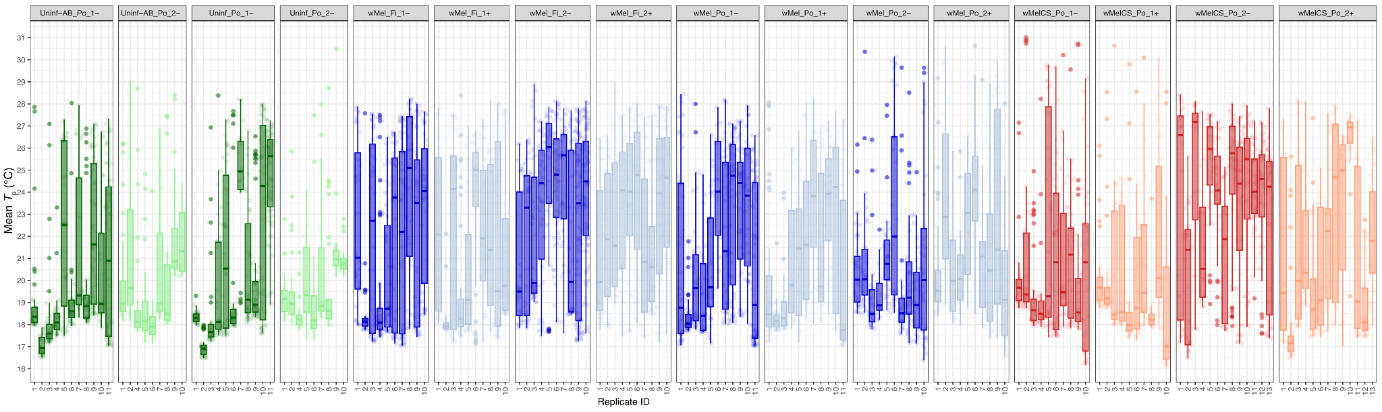

##### **Figure S2.** Similar to Figure 1, these boxplots are showing *T*_p_ in eight natural *Drosophila* strains from Finland and Portugal that were either naturally uninfected (green), or either naturally infected with wMel (blue) or wMelCS (red). The “-” symbol in the sample names and the light colors of the boxplots depict strains that underwent antibiotic treatment and subsequent restoration of the gut flora; In contrast to Figure 1, we here removed replicate experiments, where >80% of all flies cluster at the cold edge of the thermal gradient. The “***” symbol indicates a *p*-value < 0.0001.

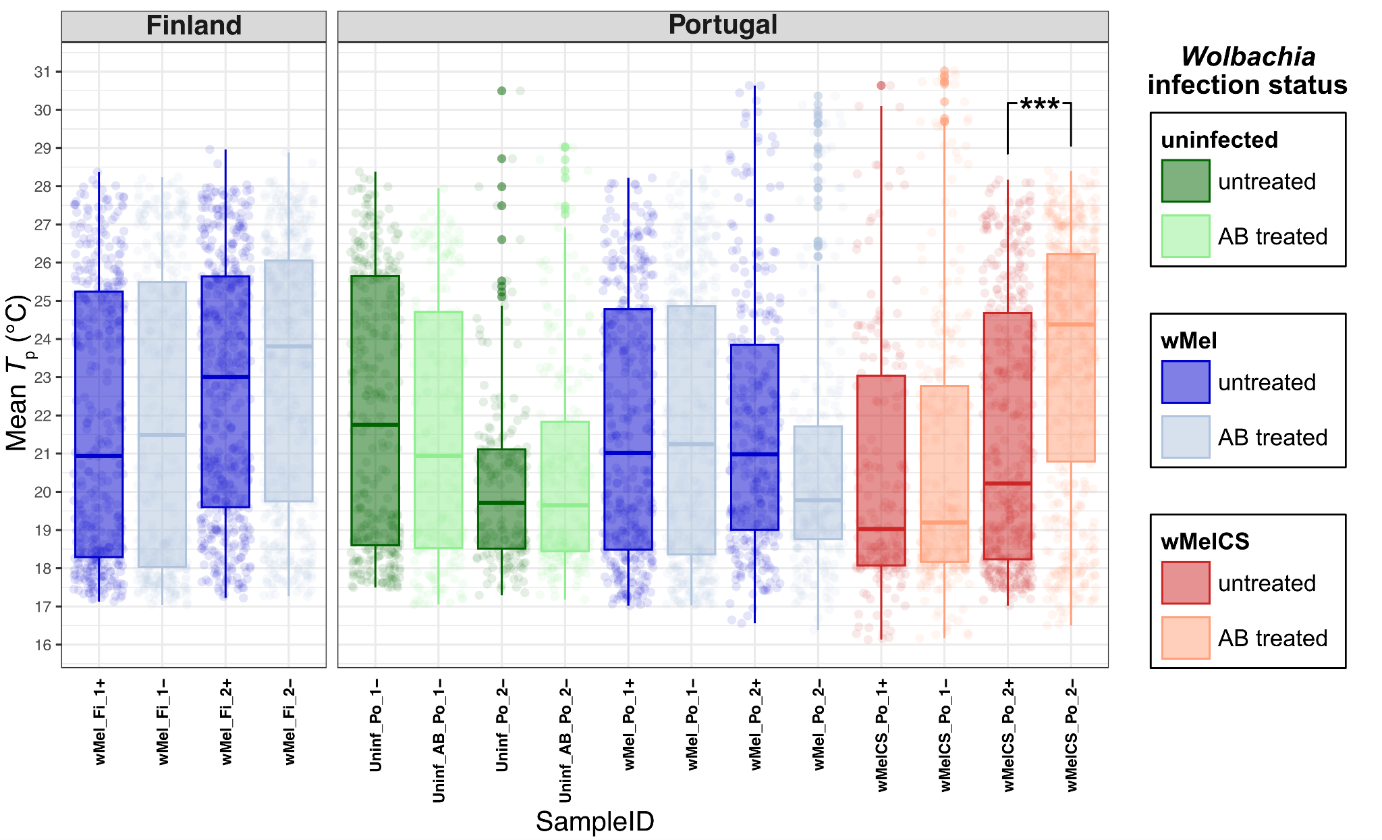
